# A Reconsideration of the Effect of Procyanidin on the Assembly of Collagen Type I

**DOI:** 10.1101/372847

**Authors:** Y. Wang, L. Jin

## Abstract

In order to elucidating the exact effect mechanism of polyphenols on the assembly of collagen, the assembled architectures of collagen treated with different amounts of procyanidin (PA) were investigated in details. The assembled morphologies of collagen were greatly influenced by the content of PA according to atomic force microcopy (AFM) images. When the content of PA was more than 20% (w/w), the fibrillar morphologies were substituted by globular aggregates, which were driven by the intense hydrogen bonding action originating from PA. While the formation of the non-fibrous aggregates was due to the coiling and entangling of flexible collagen molecules rather than their gelatinization based on the appearance of typical adsorption peaks at 222nm and 197nm on circular dichroism (CD) spectra. After being crosslinked by glutaraldehyde (GA), not only the diameters but also the lengths of fibrils increased. Unfortunately, the fibrillogenesis was still inhibited when the collagen suffered from 20% PA firstly and then 4% GA. Conversely, the fibrous morphologies of the fibrils stabilized by 4% GA and then underwent 20% PA maintained well, in spite of accompanying with grievous intertwining. This difference was derived from the change of flexibilities of collagen before and after being crosslinked by GA. Additionally, the differential scanning calorimeter (DSC) analysis confirmed the PA had no positive effect on the improvement of thermal stability of hydrous collagen, whereas the denaturation temperature of hydrated collagen stabilized by 4% GA increased from 40 °C to 80 °C.

## INTRODUCTION

In the past few decades, elucidating the mechanism of collagen assembly and controlling its aggregating architectures have achieved considerable attentions as its assembled states have profound influence on the physicochemical properties of the biomaterials (1-8). In vivo, the collagen molecules aggregate into hierarchical architectures at nano-, micro- and macro-scales, which are pivotal in serving as docking sites for some growth factors as well as maintaining the bio-functions of connective tissues (9-11). In vitro, collagen molecules assemble into various forms such as fibril, gel and sponge, respectively applied in tissue engineering scaffolds (12), drug delivery systems (13), wound dressing (14) and hemostatic (15). Evidences to date indicate that the well-organized collagen aggregates exhibit better biological characteristics than monomeric collagen molecules (16). In light of this, much researches have made on controlling the assembly of collagen molecules in vitro by adjusting the pH, ionic strength, temperature (1,8), adding polysaccharides (17) or crosslinking agents (18) in order to reconstitute collagen fibrils.

In fact, collagen molecules isolated from connective tissues exhibit poor resistance to hydrolytic and enzymatic challenges, thus the collagen materials have to be crosslinked for promoting their stability, strength and function (19,20). Surprisingly, the crosslinking action could not only promote the thermal and structural performances of collagen matrix, but also increase collagen fibrillar order and alignment (18). The usual crosslinking agents include glutaraldehyde, carbodiimide, transglutaminase, plant-derived polyphenols, and so on (19, 21).

Procyanidin (PA), well-known radical scavengers, is one of the most important classes of plant secondary metabolites, which is found in a wide variety of vegetables, flowers, fruits, seeds and bark (20-22). PA possesses a large variety of health-promoting actions, such as anti-oxidant, anti-cancer and anti-inflammatory properties (23). Besides of this, proanthocyanidins can be used as potent collagenase inhibitors and non-toxic cross-linkers for collagen matrix, especially for the bio-modification of demineralized dentin layer (24,25). Generally, the crosslinking ability of PA to collagen is attributed to its potential to induce linkages at the hierarchical architectures of collagen, while the density of the linkages significantly affects the deformation behavior of collagen (19,21,26). However, some other polyphenol compounds such as gallic acid (21), vanillic acid and syringic acid (27) evidently inhibit the self-assembly of collagen in vitro.

With these previous finding in mind, we investigated detailedly the effect of procyanidin on the assembly of collagen type I in vitro as well as the collagen fibrils stabilized by glutaraldehyde. In present study, atomic force microscopy (AFM) was applied to elucidate the assembly morphology of collagen, while circular dichroism (CD) spectrometer and differential scanning calorimeter (DSC) were used to assay the secondary structural change and thermal stability of collagen, respectively.

## MATERIALS AND METHODS

### Materials

The calf skin for isolating collagen was supplied by a local slaughter house in Shandong Province, China. The procyanidin, grape seed extract, was obtained from Tianjin Jianfeng Co. Ltd., China (more than 85% oligomeric procyanidins). The acetic acid, citric acid, Na_2_HPO_4_, NaCl and glutaraldehyde are analytical grade and purchased from Kemiou Chemical Reagent Co. Ltd., Tianjin, China. The pepsin (1:12000 units) and dialysis tube were provided by Beijing Aoboxing and Huamei Co. Ltd., China, respectively.

The acid-soluble type I collagen was extracted from calf skin according to our previous report (28). Briefly, small pieces of calf skin was digested with 5% (enzyme to substrate, w/w) pepsin in 0.5 mol/L acetic acid solution for 24 h at 4 °C in a flask equipped with a mechanical stirrer. Then the hydrolyzed product was centrifugated by GL-20G-I Refrigerated Centrifuge (Anting, Shanghai, China) at 10,000 rpm for 20 min. After that the collected supernatant was salted out by 1 mol/L NaCl solution, followed by re-dissolving of the precipitate, namely collagen type I, in 0.5 mol/L acetic acid solution. These operations were repeated for three times. Subsequently, the collagen solution was dialysed with 10 times (v/v) of 0.1 mol/L acetic acid for three days at 4 °C. Finally, a stock collagen solution of 1.0 mg/mL was stored at 4°C for use.

The buffer solutions at pH 4.0 were prepared by mixing 0.01 mol/L citric acid solution with 0.02 mol/L Na_2_HPO_4_ solution in a given volume fraction.

### The effect of procyanidin dosage on the assembly of collagen I

A given amount of the stock collagen solution was diluted in buffer solutions at pH 4.0 to obtain 12.0 μg/mL collagen solutions. Different amounts of procyanidin were added into 5 mL collagen solution above and then followed with being shaken well. The final concentrations of procyanidin were 0, 0.3, 0.6, 1.2, 2.4 and 4.8 μg/mL, i.e. the mass ratios for procyanidin to dry collagen were 0%, 2.5%, 5%, 10%, 20% and 40%, respectively. After being incubated at 20 °C for 24 h, a drop of collagen solution (about 5 μL) was placed on a new cleaved mica disc with a diameter of 6 mm, which was transferred immediately into a silicagel desiccator for drying at room temperature. The assembly morphologies of dried samples were conducted on a Dimension 3100 Nanoscope IV equipped with silicon TESP cantilevers (Shimadzu SPM-9600, Kyoto, Japan) in non-contact (taping) model at a fix scanning rate of 1Hz.

### Circular dichroism (CD) studies

The preparation of 50 μg/mL collagen solutions was similar to the method above, while the weight ratios of procyanidin to dry collagen were 0%, 10%, 20% and 40%, respectively. The mixture solutions were incubated at 20 °C for 24 h, and then the conformational changes of collagen were studied at room temperature by using Research-Grade Circular Dichroism Spectrometer (AVIV 400, Maryland, USA). All samples were scanned from 190nm to 260nm, along with a baseline correction performed with citric acid-Na_2_HPO_4_ buffer solution. The results were obtained in milli degree and converted into Molar ellipticity (deg.cm2/dmol), and corresponding graph of molar-residue-weight ellipticity versus wavelength was plotted.

### The effect of procyanidin on the assembly of collagen fibrils stabilized by glutaraldehyde

#### The effect of the amount of glutaraldehyde on the assembly of collagen I

The procedures for investigating the effect of glutaraldehyde content on the collagen assembly were totally similar to section 2.2, besides the procyanidin was replaced by glutaraldehyde. The final weight ratios of glutaraldehyde against dry collagen were 1%, 2%, 4%, and 8%, respectively. The collagen morphology was also scanned by AFM (Shimadzu SPM-9600, Kyoto, Japan) in same conditions.

#### The effect of procyanidin on the assembly of stabilized fibrils by glutaraldehyde

A given amount of glutaraldehyde was add into 12.0 μg/mL collagen solution as experiment above, and the concentration of GA was 0.48 μg/mL. After being incubated at 20 °C for 24 h, 20% of procyanidin was added and followed by the other 24 h incubation at 20 °C. As a control, similar experiments were carried out except that the order of adding glutaraldehyde and procyanidin was changed. Finally, the aggregated architectures of collagen were observed by using SPM-9600 (Shimadzu, Kyoto, Japan).

### Differential scanning calorimeter (DSC) measurements

The collagen samples were prepared as follow. 0%, 5%, 10% and 20% by weight of procyanidin was added into 1 mg/mL collagen solution respectively, and then these solutions were incubated at 20 °C for 24 h to be tested. Additionally, 20% of procyanidin was added into 1mg/mL collagen solution for incubation at 20 °C, after 24 h 4% glutaraldehyde was placed into the solution above which was successively incubated in same condition. As a contrast, the glutaraldehyde reacted with collagen firstly and then procyanidin was used, and other experimental factors were unchanged.

To demonstrate the influence of individual procyanidin or glutaraldehyde and their synergism on the stability of collagen, DSC analysis was conducted on DSC 200 PC (Netzsch, Selb, Germany) in the temperature ranged from 20 °C to 100 °C with a constant heating rate of 5 °C/min under a nitrogen flow. About 10 mg collagen solution mentioned above was sealed in an aluminum pan with a sealed aluminum pan containing 10 mg buffer solution as the reference.

## RESULTS AND DISCUSSION

### The effect of procyanidin dosage on the assembly of collagen I

The assembly morphologyies of collagen type I in the presence of different contents of procyanidin were shown in Figure 1a-f. It can be found that the procyanidin affected pronouncedly on the assembly behavior of type I collagen as the dosage of procyanidin increasing from 0% to 40%. Without procyanidin, the self-assembled collagen fibrils with diameters less than 3 nm almost paralleled to each other, shown as the arrows in Figure 1a. Distinctly, the lengths of the collagen fibrils were much more than 300nm, which indicated that the collagen molecules self-assembled longitudinally while the lateral assembly was inhibited significantly. It was because that the amino groups on side chain of collagen molecules were positively charged at pH 4, and the electrostatic repulsion among collagen molecules prohibited the lateral aggregation of collagen (8). When 2.5% procyanidin reacted with collagen, the orientation of collagen fibrils was disturbed slightly and a mesh-like architecture was obtained, similar to previous report (19). In this case, the limited amount of procyanidin molecules distributed longitudinally along the collagen fibrils and hydrogen bonds had formed among procyanidin and collagen fibrils. As the content of procyanidin increased to 5%, the hydrogen bonding action became stronger and some bulges on the cross-points of fibrillar network were observed clearly (shown in Figure 1c). When the content of procyanidin was at 10%, these bulges became more and bigger while some disordered fibrils arranged randomly throughout the bulges. This was probably attributed to the changing of fibrous conformation of collagen under strong hydrogen bonds provided by procyanidin. In addition, some charged amino groups on collagen molecules might be shielded by procyanidin, thus the electrostatic repulsion of charged collagen molecules was decreased heavily. Even worse, the fibrillation of collagen was inhibited almost completely when the dosage of procyanidin was much than 20%, shown in Figure 1e and 1f. Instead of collagen fibrils, some globular collagen aggregates appeared. Especially in the case of 40% procyanidin, no collagen fibril was observed while the diameter of these globular aggregates decreased. A reasonable explanation for this phenomenon was that a large amount of procyanidin molecules distributed longitudinally on collagen and aggregated together by intense hydrogen bonds, and thus the collagen molecules coiled randomly to form some globules. The more the content of procyanidin, the more the number of elliptic aggregates, hereby their diameters became much less (shown in Figure 1f). Collectively, the assembly behavior of collagen influenced by procyanidin was a procedure driven by hydrogen bonds, in which the intensity of the hydrogen bonds was a crucial factor. Besides of this, the large length-diameter ratio (300:1.5) and flexibility of collagen molecule was in favor of the entanglement.

**Figure 1.**
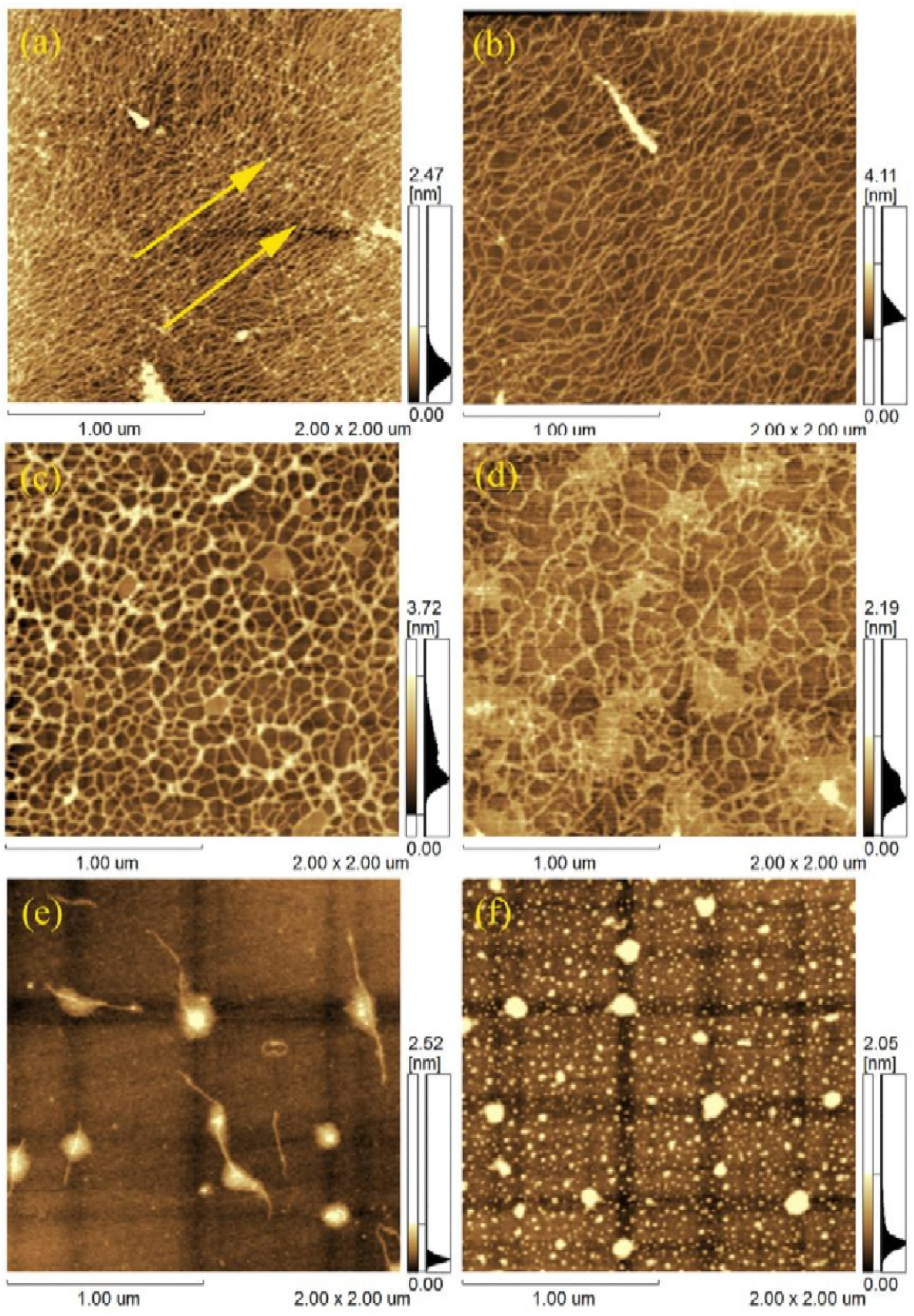
AFM images of the assembly of type I collagen affected by different dosage of procyanidin: (a) 0%, (b) 2.5%, (c) 5%, (d) 10%, (e) 20% and (f) 40%.

Generally, the stability of fibrils self-assembled from collagen molecules in vitro depended mainly on intermolecular hydrogen bonds. While the procyanidins are condensed by monomeric flavan-3-ol unites which are rich in phenolic hydroxyls, and thus this hydrogen-bond interaction would be strengthened as the increasing of procyanidin content. However, the ordered hydrogen bonding was disturbed when the concentration of procyanidin was too high because the collagen molecules were completely encapsulated by procyanidin molecules, and consequently the fibrous assembly was imbalanced. It has been reported that the fibrillogenesis of collagen was inhibited by the presence of natural polyols, such as glucose, fructose, methylglucoside and sorbitol by disturbing the hydrogen bond interactions between collagen helices (29). According to our research, the inhibition was probably due to the change of the fibrillar conformation of collagen caused by strong hydrogen bonding, which was disadvantage for the fibrillogenesis.

### Circular dichroism (CD) studies

In order to elucidate the mechanism of the effect of procyanidin on the assembly of collagen, the secondary structure of collagen treated with procyanidin was investigated by CD. The CD spectra of type I collagen treated by different concentration of procyanidin were shown in Figure 2. For the case of natural type I collagen, specific peaks in far ultraviolet (UV) region at 222nm and 197nm can be observed on CD spectrum, which were due to n to π* transition of polyproline II ring and π to π* transition, respectively (27, 30-32). Shown as Figure 2, the absorption bands on CD spectrum (curve a) were typical peaks of collagen type I, which indicated that the secondary structure of collagen kept well. Compared with native collagen, the peaks at 222nm of collagen treated by different amount of procyanidin had little change, while the peak intensities at 197nm decreased slightly as the increase of procyanidin concentration. It is confirmed that a red shift in 222nm signal would occur when the collagen denatured or its triple helical structure unfolded (27). However, there was no red shift was observed on the spectrum of collagen treated by procyanidin.

**Figure 2.**
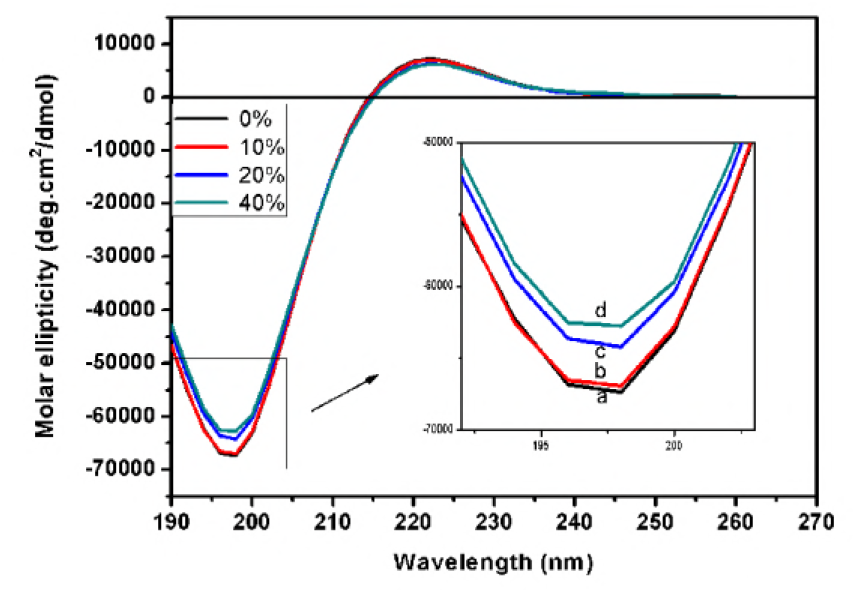
Circular dichroism spectra of type I collagen treated by different concentration of procyanidin: (a) 0%, (b) 10%, (c) 20% and (d) 40%. The inset was an enlarged spectra at 190-203nm.

Moreover, the corresponding molar ellipticity and Rpn (the ratio of positive peak intensity to negative peak intensity) values were listed in Table 1. The Rpn value of type I collagen was 0.107, and the Rpn value was 0.106 in the case of adding 10% of procyanidin. And the Rpn value decreased to 0.098 when the dosage of proyanidin increased to 40%. It has been reported that the micro unfolding of the collagen triple helical structure rather than denaturing was observed under the effect of phenolic compound and the corresponding Rpn values decreased (27, 33-34). Therefore, the decreasing of Rpn values here was probably attributed to the conformational change of collagen molecule, which agreed well with the investigation of AFM (globular aggregates shown in Figure 1e and f). Thus, it could be concluded that the procyanidin had not leaded to the change of collagen secondary structure according to CD results.

**Table 1.** Molar ellipticity and Rpn (the ratio of positive peak intensity to negative peak intensity) values of collagen treated with different amount of proanthocyanidin

| Dosage of procyanidin<br>(%) | Molar ellipticity (deg.cm<sup>2</sup>/dmol) |  | Rpn |
| --- | --- | --- | --- |
|  | 220(nm) | 197(nm) |  |
| 0 | 7184.9 | -67353.3 | 0.107 |
| 10 | 7071.0 | -66909.2 | 0.106 |
| 20 | 6325.6 | -64197.3 | 0.098 |
| 40 | 6201.2 | -62744.7 | 0.098 |

In fact, the phenolic hydroxyl groups on procyanidin can cross-link with collagen by hydrogen bonds, and consequently the stability of collagen was increased. Therefore, the secondary structure of collagen was not destroyed by procyanidin, which agreed with the previous study elsewhere (19). However, in the case of acid soluble collagen, the presence of a large amount of procyanidin had negative influence on the fibre forming from collagen molecules. It was probably because of the change of conformation of collagen under the intense hydrogen bond action of procyanidin rather than the denaturation of secondary structure of colllagen. It was to say the formation of the oval aggregates was not gelatinizing of collagen.

### The effect of procyanidin on the assembly of collagen fibrils stabilized by glutaraldehyde

#### The effect of glutaraldehyde on the assembly of collagen I

The effect of the dosage of glutaraldehyde on the assembly of collagen was shown in Figure 3. Obviously, the diameters of collagen fibrils increased distinctly as the increase of glutaraldehyde dosage. Compared with the collagen fibrils without being crosslinked by glutaraldehyde (shown as Figure 1a), the fibrillar morphology changed slightly when 1% glutaraldehyde was used (shown as Figure 3a). After being treated by 2% glutaraldehyde, the lateral crosslinking among collagen fibrils became clearly and some collagen fibrils with thicker diameter appeared, which was shown in Figure 3b. When the dosage of glutaraldehyde was above 4%, branched collagen fibrils with length of micrometers were observed. In fact, glutaraldehyde is a common crosslinker of biomaterials to improve their stability against heat or enzyme (35-38) depending on the reaction of aldehyde group with amino group on side chain of collagen molecule. Thus the collagen fibrils aggregated closely through intense lateral assembly, shown as Figure 3d. Thanks to the crosslinking action of glutaraldehyde, not only the diameter but also length of collagen fibrils was improved greatly.

**Figure 3.**
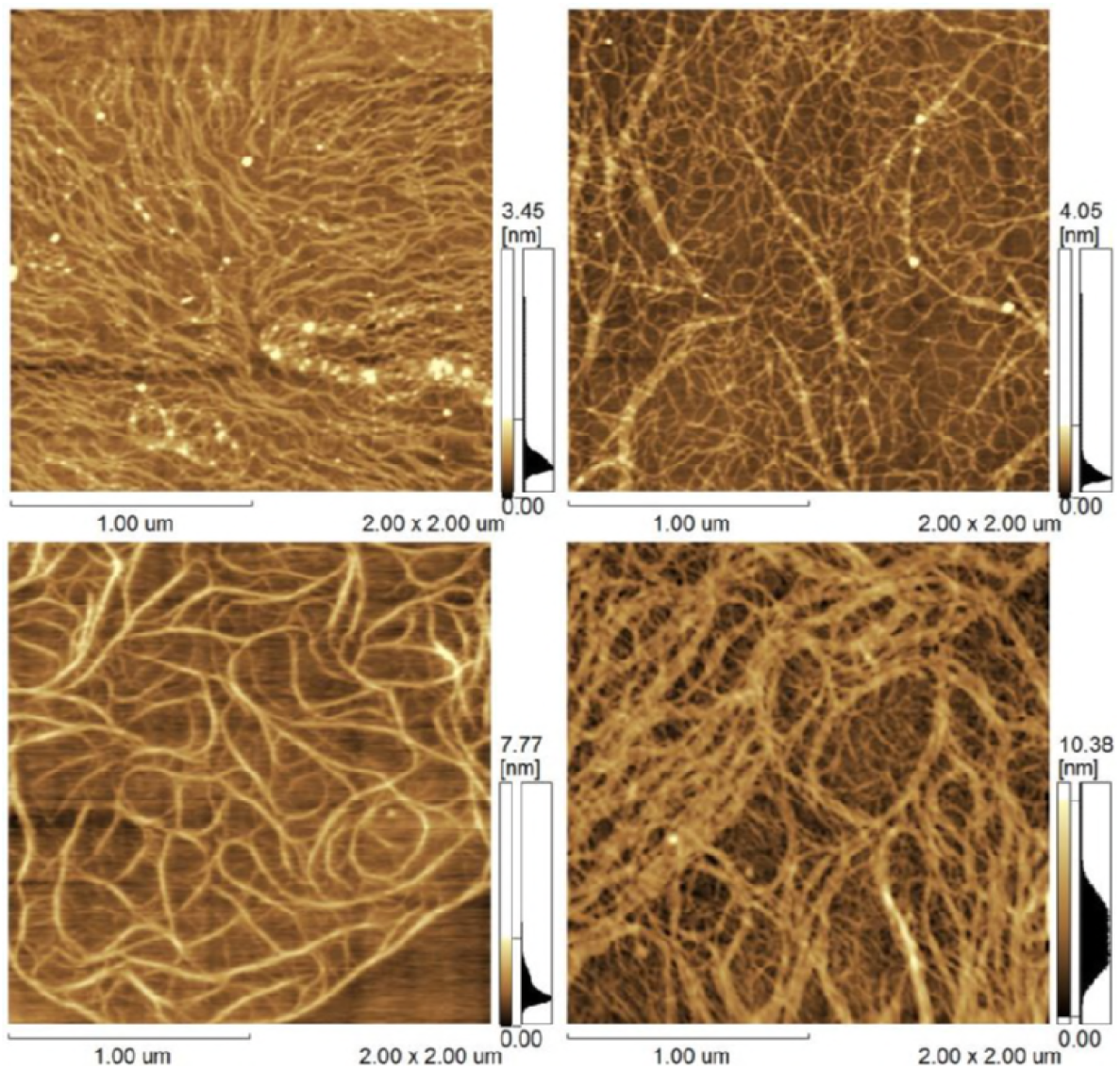
The effect of glutaraldehyde on the assembly of collagen type I at pH 4: (a) 1%, (b) 2%, (c) 4% and (d) 8%.

#### The effect of procyanidin on the assembly of stabilized fibrils by glutraldehyde

The assemble morphology of collagen which was treated by 20% procyanindin firstly and then by 4% glutaraldehyde was shown in Figure 4a. Some irregular aggregates, which interlaced with several fibrils, were observed as the crosslinking action of glutaraldehyde. Whereas in the case of 4% glutaraldehyde which was followed by 20% procyanidin, collagen fibrils were recorded clearly only that these fibrils coiled and tangled with each other. For the former, the conformation of collagen changed from extended chain to random coil on account of the strong hydrogen bonding action to form compact micro particles. Compared with the former, covalent bonding actions were constructed among the collagen molecules depending on the crosslinkage of glutaraldehyde and thus stable collagen fibrils formed. It should be note that the slenderness ratios and the flexibilities of the fibrils were also high, which was in favor of their coiling and tangling with each other. As the adding of excess procyanidin, a large amount of hydrogen bonds inter-fibrils and intra-fibrils formed, consequently these flexible fibrils folded, coiled and twined, shown as figure 4b.

**Figure 4.**
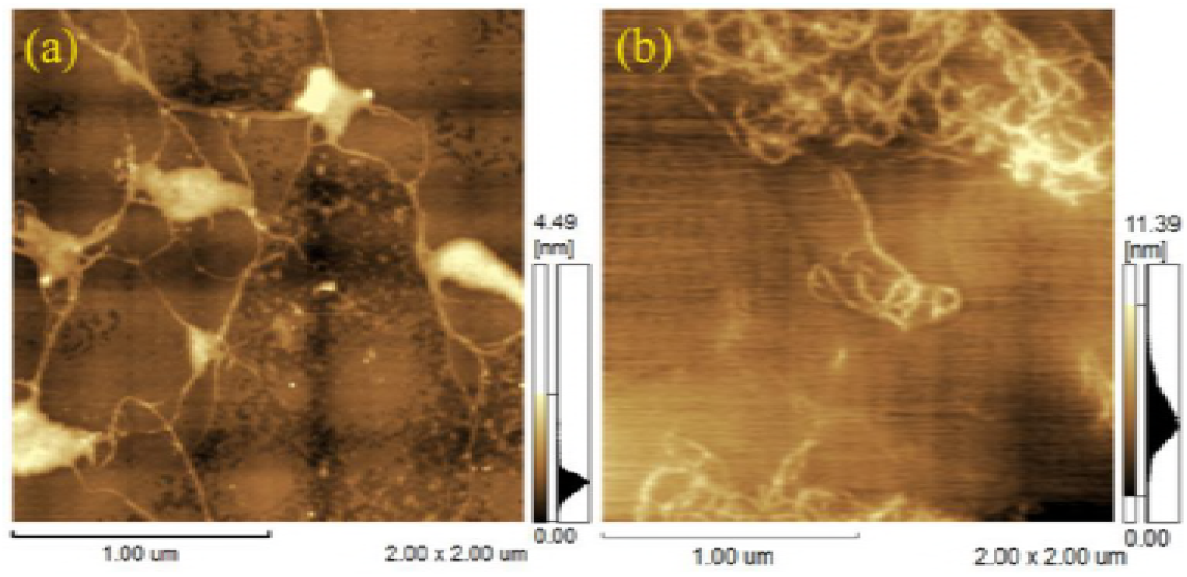
The assembly architectures of collagen treated by: (a) procyanidin followed with glutaraldehyde and (b) glutaraldehyde followed by procyanidin.

After being crosslinked by glutaraldehyde, the stability and stiffness of collagen fibrils were increased greatly, so limited intertangling taken place when some external forces put upon these fibrils. In spite of this, the fibrillar topographies were also kept well. However, the stability of self-assembled collagen fibrils was much less than that of crosslinked fibrils, and thus the fibrillar morphology could not be maintained when the collagen were treated by superfluous procyanidin. In another word, the assemble morphology of collagen influenced by procyanidin was depending greatly on the structural stability of collagen fibrils.

### The Thermal Stability of Collagen Investigated by DSC

DSC was a useful instrument for investigation the improvement of chemicals to collagen. So the thermal stability of collagen treated with different amount of procyanidin was investigated by DSC, and their DSC curves were shown in Figure 5. On the DSC thermograph of collagen (Figure 5a), an endothermic peak at about 40 °C was detected, which was approximate to the value reported by previous literature (39). It is known that this peak is considered to be associated with the denaturation of native structure of collagen type I, which was often accompanied with the shrinkage of collagen (40,41). When the dosage of procyanidin was 5%, a similar peak at 40 °C was recorded, shown as Figure 5b. As the content of procyanidin increased to 10%, the endothermic peak at 40 °C became weaker. All these peaks were attributed the endothermic helix to coil transition (42). However, the endothermic peak was un-conspicuous when 20% of procyanidin was used to treat collagen. It was probably because that the long rod-like collagen molecules had coiled and tightly agglomerated with each other, hence the shrinking and denaturating of collagen was unnoticeable or difficult to undergo at the same temperature.

**Figure 5.**
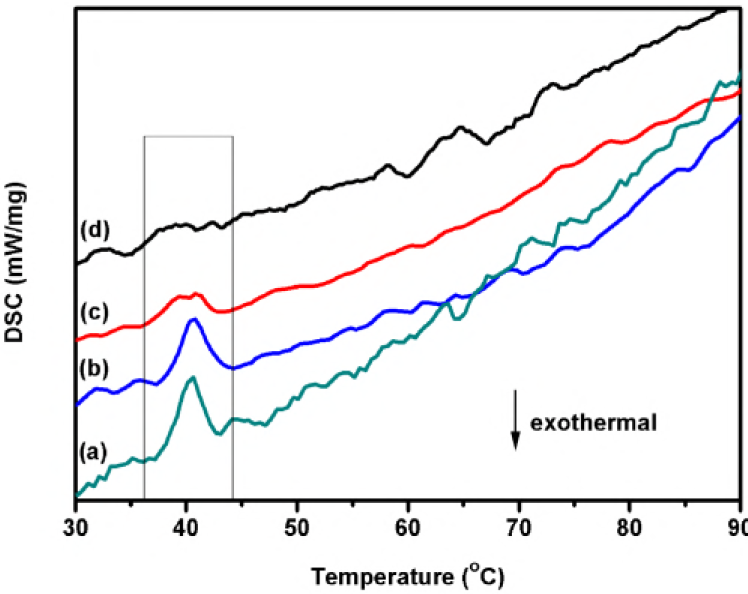
The differential scanning calorimeter (DSC) curves of collagen treated by using procyanidin: (a) 0%, (b) 5%, (c) 10% and (d) 20%.

Furthermore, the effects of procyanidin integrated with glutaraldehyde (marked as PA-GA), glutaraldehyde as well as glutaraldehyde incorporated with procyanndin (identified as GA-PA) on the collagen stability were researched, respectively. As shown in Figure 6a, the denaturation temperature (T_d_) of collagen treated by glutaraldehyde was increased to 80 °C, which reflected the thermal stability of collagen was improved greatly. The T_d_ of collagen treated with GA-PA was approximate to it of collagen crosslinked only with glutaraldehyde, and the peak value was at around 80 °C, shown as Figure 6b. Differently, the width of the peak become wider which was attributed to the orientation of the collagen fibrils was disturbed (43), shown as Figure 4b. Compared with the DSC curves above, no obvious endothermic peak was measured in the case of collagen treated with PA-GA, which was similar to it of collagen treated with procyanidin only and shown in Figure 6c. It was probably because that the aggregated complexes of coiled collagen with PA-GA had been hard to shrink or deform at this temperature.

**Figure 6.**
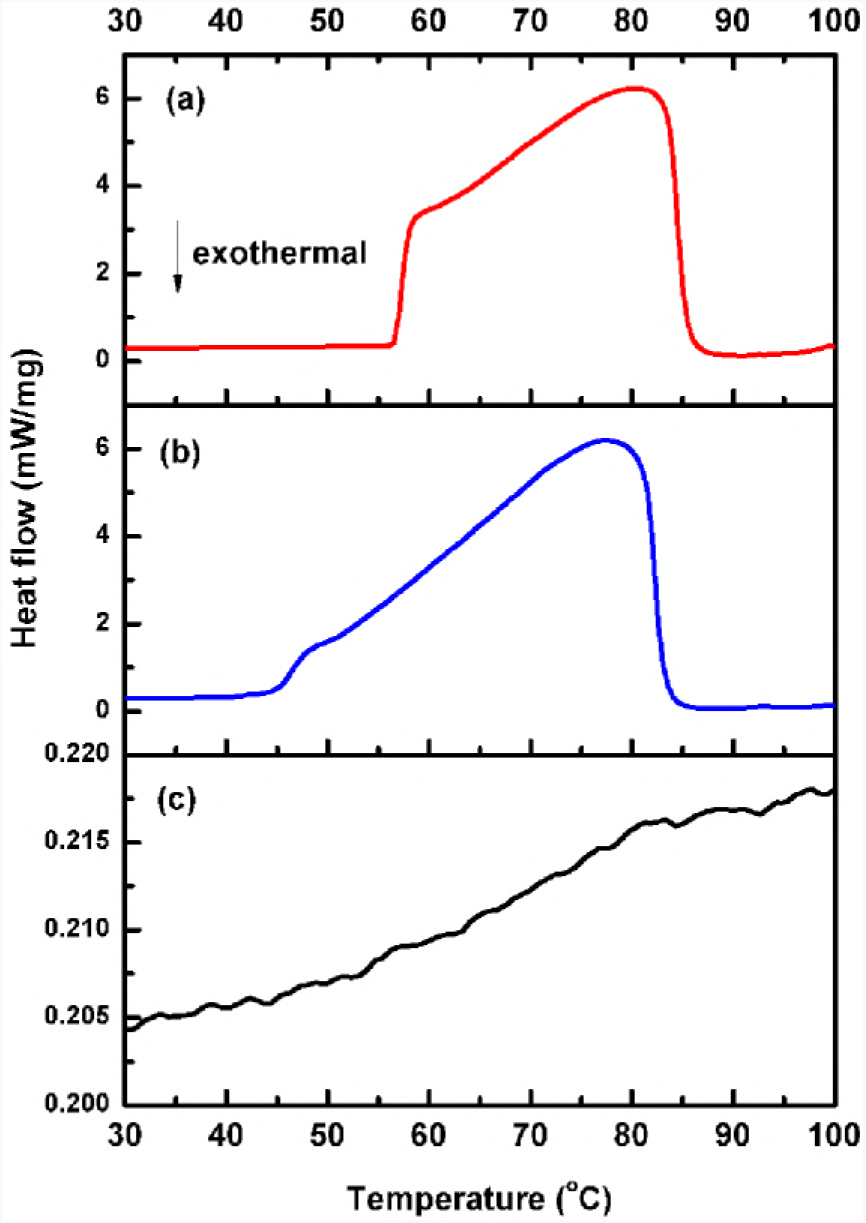
The differential scanning calorimeter (DSC) curves of collagen treated by: (a) glutaraldehyde, (b) glutaraldehyde followed by procyanidin and (c) procyanidin followed by glutaraldehyde.

It was reported that the shrinkage temperatures (denaturation temperature) of hide powder treated by condensed tannins was about 80 °C as the stabilization of the vegetable polyphenols through hydrogen bonds. And it increased to around 107 °C when the hide powder was crosslinked by condensed tannins combined with aldehydic agent. It was owing to the formation of new covalent bonding among collagen and phlyphenols by the crosslinking of aldehydic agent, which was known as synthetic effect of tannin-aldehyde (44-46). Unfortunately, no thermal stability improving was observed when the collagen was treated by procyanidin individually. Besides of that, the synthetic effect of tannin-aldehyde was not remarkable in our experiments. Possibly, this divergence was resulted from the structural difference between acid soluble collagen molecule and native collagen fibrils of connective tissues.

As is well known, the triple helical collagen molecules self-assemble into the characteristic axial D-periodicity to form microfibrils and fibrils in vivo, in which the inter-collagen covalent cross-links between lysine or allysine residues at the overlap position of collagen molecules are crucial for the formation of 4D-staggered arrangement (47-48). Importantly, these intermolecular cross-links are a prerequisite for the physical and mechanical performances of collagen fibrils (^48^). However, these lateral cross-links were broken as the partly cleavage of telopeptides by pepsin in the extraction procedure of collagen. And no new covalent crosslinking formed in the self-assembled collagen fibrils, even though procyanidin had been used. As a consequence, the thermal stability of collagen had not been improved by the procyanidin as the self-assembled fibril itself was lack of covalent crosslinks. Conversely, the collagen molecules were covalently cross-linked by the glutaraldehyde, which was similar to the natural cross-links in native tissues. And the significant stabilization of glutaraldehyde was reflected well in the results of DSC thermograph and AFM imagines shown above.

### The assembly mechanism of collagen in the presence of procyanidin and glutaraldehyde

On the base of the analysis aforementioned, a schematic illustration about the effect of procyanidin on the aggregating of the self-assembled collagen fibrils and stabilized fibrils by glutaraldehyde was shown in Figure 7. The influence of procyanidin on the assembly of collagen was closely related to the dosage of procyanidin. When the concentration of procyanidin was low, the procyanidin molecules longitudinally distributed on the self-assembled collagen fibrils by hydrogen bonds. As the increasing of the procyanidin content, the orientation of collagen fibrils was disturbed and fibrils were inclined to coil, thereby a mesh-like morphology appeared. Suffered from the strong hydrogen bonding action of a large amount of procyanidin, the collagen fibrils coiled and entangled into irregular aggregates and few of fibrils could be observed, even though the second structure of collagen was not destroyed. In this process, both the flexibility of collagen and the absence of covalent crosslinkages inter-collagen molecules were benefit for the conformational changes and the formation of globular morphology, while the intense hydrogen bonds were the driving forces. In the case of collagen fibrils stabilized by glutaraldehyde, lateral crosslinking action among collagen molecules was constructed by the reaction of amino groups on side chain of collagen with aldehyde groups, and as a result the stiffness and the stability of the fibrils was improved greatly. Therefore, the cross-linked fibrils maintained the fibrous morphology well in spite of folding and twisting with each other in the presence of an abundant of procyanidin. The intermolecular crosslinking from glutaraldehyde played an important part in keeping the fibrillar conformation of collagen, which was comparable to the important function of the cross-links between lysine or allysine residues on adjacent collagen molecules.

**Figure 7.**
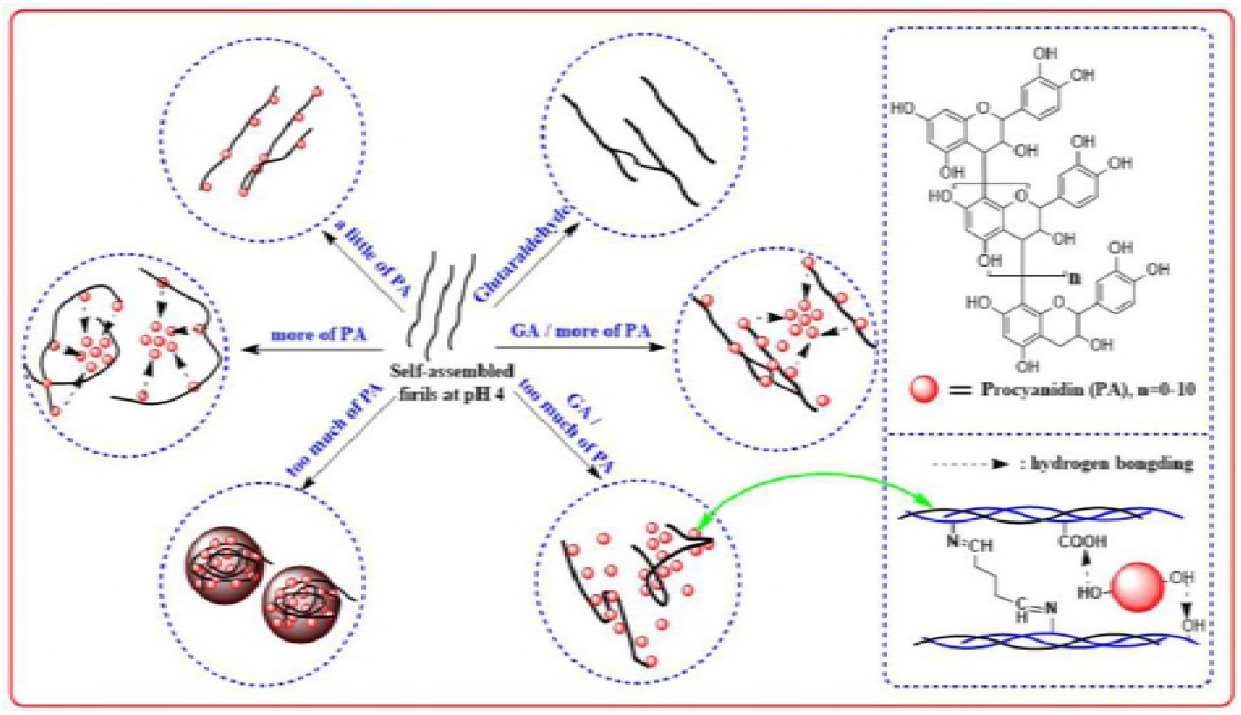
Schematic illustration about the effects of procyanidin on the aggregating of self-assembled collagen fibrils and stabilized fibrils by glutaraldehyde.

It was noteworthy that this research was not to deny the important role of procyanidin in improving the stability of collagen but to emphasis the significance of the intermolecular covalent crosslinks like in vivo in maintaining the fibrillar conformation of acid soluble collagen. Furthermore, the improvement of procyanidin to collagen was frequently based on the intermolecular covalent cross-linking so as to enhance the physical and mechanical performances of assembled collagen fibrils. And more intuitive evidences were supplied here.

## CONCLUSIONS

The effect of procyanidin on the assembly of collagen in vitro was investigated in details, and it was found that this influence was related closely with the dosage of procyanidin. When the content of procyanidin was less than 10% (to dried collagen, w/w), the fibrillar morphologies of self-assembled collagen fibrils would keep well. However, the collagen would aggregate into irregular and tight globules when the content of procyanidin was more than 20%, while the diameters of the globules decreased as the increasing of procyanidin content. It was attributed to the conformation change driven by intense hydrogen bonding action provided by procyanidin rather than denaturation of secondary structure of collagen, in which the high slenderness ratio and flexibility of collagen as well as the absence of covalent crosslinkages inter-collagen molecules were benefit for the conformational changes and the formation of globular architectures. If the self-assembled collagen fibrils were modified by cross-linker such as glutaraldehyde, not only the thermal stability but also the conformational rigidity of the fibrils were improved greatly, thus the fibrous morphologies would be maintained besides of some coiling and entangling when excess procyanidin was used. These results demonstrated visually the significance of the intermolecular covalent crosslinks in maintaining the fibrillar conformation of acid soluble collagen in vitro.

## ACKNOWLEDGMENT

This paper was supported by the National Key Research and Development Program of China (2017YFB0308503) and the Shandong Provincial Key Research and Development Program, China (2017GGX80103). The authors thanks for the guidance of Professor Bi Shi from Sichuan University and the conveniences supplied by National Engineering Laboratory for Clean Technology of Leather Manufacture in measurement.

## Author Contributions

Y. Wang designed and performed all the experiments, analyzed the data and wrote the paper. L. Jin took part in some discussion. All authors have given approval to the final version of the manuscript.

## Notes

The authors declare no competing financial interest.

